# Low beta power reflects perceived temporal probabilities for uncertain future events

**DOI:** 10.1101/234070

**Authors:** Alessandro Tavano, Erich Schröger, Sonja A. Kotz

**Affiliations:** BioCog, Cognitive Incl. Biological Psychology, Institute of Psychology, University of Leipzig, Leipzig - 04109, Germany; Department of Neuroscience, Max Planck Institute for Empirical Aesthetics, Frankfurt am Main – 60322, Germany; Department of Neuropsychology, Max-Planck-Institute for Human Cognitive and Brain Sciences, Leipzig - 04103, Germany; Faculty of Psychology and Neuroscience, Department of Neuropsychology and Psychopharmacology, Maastricht University, Maastricht - 6200 MD, The Netherlands.

## Abstract

Humans tend to use elapsed time to increase the perceived probability that an impending event – e.g., the Go sign at a traffic light - will occur soon. This prompts faster reactions for longer waiting times (hazard rate effect). Which neural processes reflect instead the perceived probability of uncertain future events? We recorded behavioral and electroencephalographic (EEG) data while participants detected a target tone, rarely appearing at one of three successive positions of a repeating five-tone sequence with equal probability. Pre-stimulus oscillatory power in the low beta-band range (Beta 1: 15-19 Hz) predicted the hazard rate of response times to the uncertain target, suggesting it encodes abstract estimates of a potential event onset. Informing participants about the target’s equiprobable distribution endogenously suppressed the hazard rate of response times. Beta 1 power still predicted behavior, validating its role in contextually estimating temporal probabilities for uncertain future events.

**Highlights:**

- Elapsed time to an uncertain future target increases response speed (Hazard rate).
- Pre-stimulus low beta-band (Beta 1: 15-19 Hz) power predicts the hazard rate to uncertain targets.
- Beta 1 power predicts response times even when elapsed time is factored out.

**eTOC Blurb:** Tavano et al. show that pre-stimulus low beta band (15-19 Hz) power predicts response times to an uncertain future target, even before its occurrence and under different prior knowledge conditions, suggesting it reflects contextual, subjective estimates of potential future events.

## Introduction

How do humans successfully anticipate when a target event will occur within a given time window? One way is to keep track of the elapsed time to a target that must appear within a specific interval and to use it to incrementally increase attention, thus gaining in accuracy and response speed (Bermudez and Schultz, 2014; Bertelson, 1967; Bolger et al., 2014; Correa et al., 2004, 2006; Correa and Nobre, 2008; Coull, 2009; Coull et al., 2011; Coull and Nobre, 2008; Cui et al., 2009; Griffin et al., 2001; Janssen and Shadlen, 2005; Jones et al., 2002; Luce, 1986; Meck, 1988; Nobre and Coull, 2010; Niemi and Näätänen, 1981; Vangkilde et al., 2012; Woodrow, 1914). The time-dependent performance gain for such a deterministic target is underlain by an increase in perceived event probability, termed the *hazard rate* of events. The hazard rate is generated by normalizing the implicit estimates of target probability at any point within the time window (*probability density*) by the ever-diminishing probability that the target has not yet occurred (*survival probability*, Elithorn and Lawrence, 1955; Luce, 1986).

Neural circuits are sensitive to the hazard rate of deterministic targets. The groundbreaking work of Janssen and Shadlen (2005) showed that neurons in the macaque Lateral Intraparietal Area (LIP) maintain a neocortical representation of the hazard rate. Firing rates in LIP neurons increase with elapsed time irrespective of whether a motor response is required (Maimon and Assad, 2006; Yang and Shadlen, 2007; for a similar result in monkey prefrontal cortex, see Genovesio et al., 2006). In humans, hazard rate effects on cortical activity have been found in regions homologous to the LIP area (Intraparietal Sulcus, IPS: Cotti et al., 2011; Coull et al., 2014; Davranche et al., 2011; Inferior Parietal Cortex, IPC, Bolger et al., 2014), but also in motor, sensory, and prefrontal regions (Supplementary Motor Area, SMA, and the right Superior Temporal Gyrus, STG, for auditory stimuli, Cui et al., 2009; premotor cortex, Hultin et al., 1996; the right Prefrontal Cortex, PFC, Coull, 2009), suggesting the existence of a distributed brain network for the hazard rate (Coull et al., 2011).

However, real-life events are seldom deterministic. We asked which neural processes underlie the anticipation of target events that may or may not occur within a given time window. The uncertainty about the onset of a target event should force participants to entertain abstract estimates of event probability. We collected human behavioral and electroencephalographic (EEG) data while participants reacted to low-pitch target tones (349 Hz) appearing only 20% of the times within a continuously repeating set of four standard tones (440 Hz) followed by higher-pitch deviant one (494 Hz), indicating the end of a sequence (Sussman et al., 2002; Sussman and Gumenyuk, 2005).A target could appear at either standard tone position two, three, or four with equal probability (Figure 1A and Experimental procedures). Elapsed time was represented by the isochronous unfolding of potential target positions within each repeating tone sequence. We expected participants to iteratively build up and update target probability estimates within each sequential cycle so that response times would reflect a hazard rate distribution (Figure 1B). We hypothesized that perceived temporal probability estimates would change the weights of post-stimulus sensory processing, modulating an early, sensory-specific prediction error reflected by a deviant N1 response between 100 and 150 ms post-stimulus onset (Friston, 2005; Garrido et al., 2009a,b; Jaramillo and Zador, 2011; Tavano et al., 2014; Wacongne et al., 2012) as well as a late target-related prediction error reflected by the N2b and P300 responses peaking at ~ 200 ms and ~ 300 ms (Ahveninen et al., 2002; Chennu et al., 2013; Miniussi et al., 2001; Nieuwenhuis and Yeung., 2005; Nieuwenhuis et al., 2011; Schröger and Wolff, 1998; Schröger et al., 2015; Sokolov et al., 2002; Sussman, 2007).

**Figure 1.**
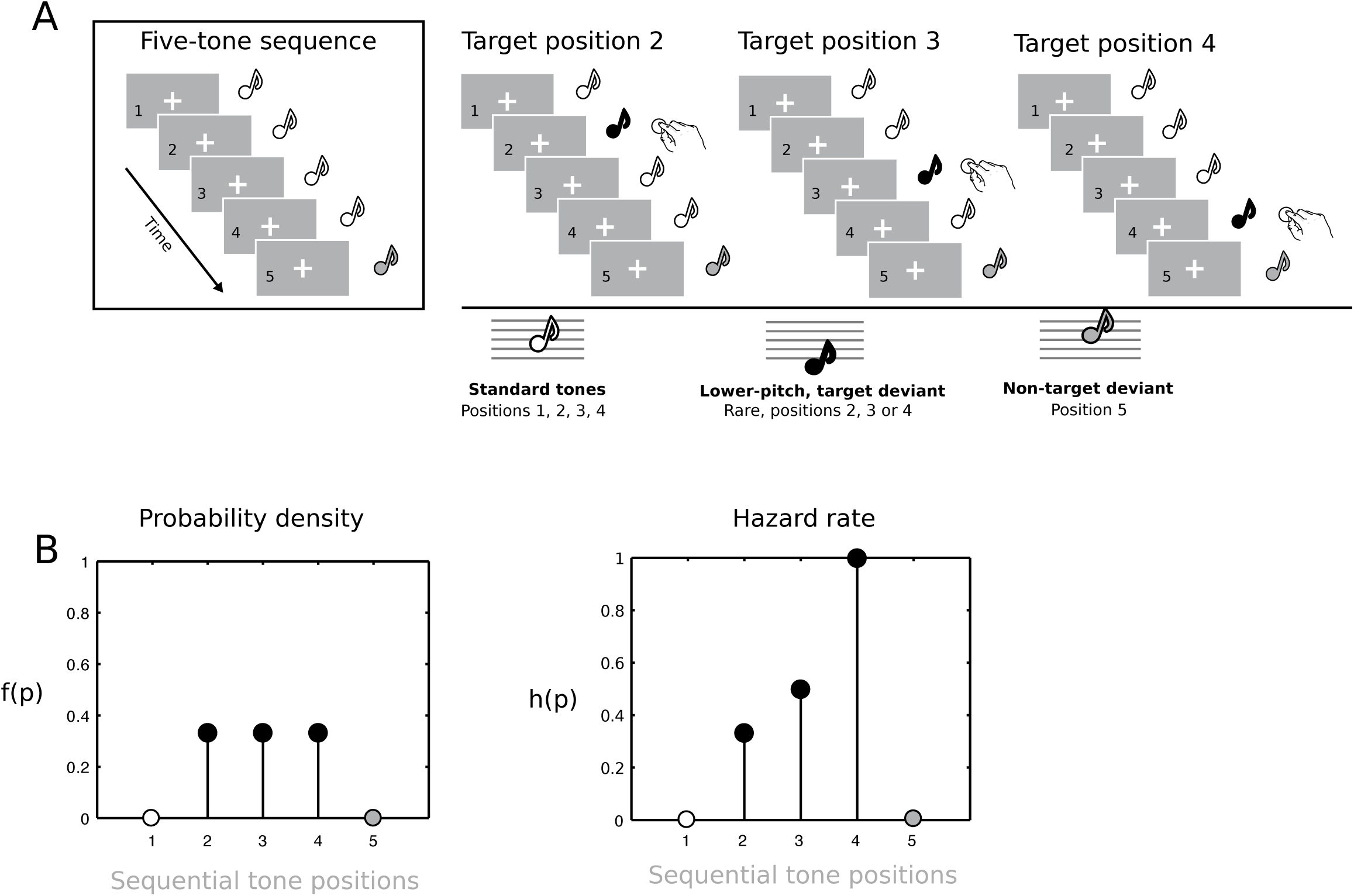
A: Frequently repeating five-tone sequences (left panel, four A_4_ tones followed by a B_4_ tone) were rarely (20%) interspersed with sequences containing a target deviant tone (F_4_) at either of three equally probable positions (position 2, 3, and 4, right panel). B: Left, the discrete probability density (all potential target onset positions have the same probability within each condition) representing the actual target probabilities. Right, the resulting hazard rate model. The first standard tone and the last non-target deviant tone never host a target.

However, if an internal representation of potential target probability determines response speed, we should observe rapid probability update processes taking place ahead of a target’s onset. Recent work suggests that pre-stimulus activity modulation in the beta band (13-30 Hz) tracks the temporal regularity of isochronous tone sequences (Fujioka et al., 2012, 2015; Merchant et al., 2015) and correlates behavioral accuracy in detecting temporally irregular stimulus delivery (Arnal and Giraud, 2012; Arnal et al., 2015; Patel and Iversen, 2014). In purely perceptual tasks, beta-band power decreases after tone onset and then sharply rebounds (event-related synchronization, ERS, Pfurtscheller et al., 2003) when approaching an immediately following tone (Fujioka et al., 2012, 2015). We hypothesized that the single-trial dynamics of beta-band oscillatory power may extend beyond a rhythm tracking function and address the fundamental question of how humans internally represent the accrual of probability estimates in time. Engel and Fries (2010) first framed a role for beta-band oscillations in endogenously encoding the intended or predicted maintenance of a current internal state, may it be cognitive or motor. Cortical feedback connections projecting top-down information tune into the beta band (Michalareas et al., 2016; Lee et al., 2013). Spitzer and Haegens (2017) recently expanded on the content-specific function of endogenous synchronization in the beta band by suggesting that beta oscillations create short-lived neural assemblies, which help to reactivate task-relevant information. We specifically predicted that hazard rate effects would be encoded in the low beta-band range or Beta 1 rhythm (< 20 Hz), following converging evidence from three independent lines of work: 1) Simulation work on neural activity across cortical layers suggests a role for Beta 1 oscillations in distinguishing between novel and standard events (Kopell et al., 2011); 2) Motor output inhibition has been more frequently linked to high beta-band or Beta 2 activity (> 20 Hz) (Brovelli et al., 2004); 3) Low beta-band (~15 Hz) activity of basal ganglia origin appears to encode the evaluation of an event’s task relevance (target vs. non-target) regardless of motor output (Leventhal et al., 2012).

The experiment was organized into two sessions. In a first session, participants were uninformed about the target’s distribution: neural (sensory prediction error and Beta 1 oscillations) and behavioral measures should display a hazard rate distribution. In a second session, we probed whether providing prior information on the target’s uniform distribution would suppress the hazard rate and its neural correlates.

## Results

Behavioral and EEG data were recorded as participants (N = 26) completed the experiment, organized as a 2 × 2 design with factors *Prior Information* and *Stimulus Onset Asynchrony* (SOA). The factor Prior Information tested the effect of providing participants with explicit information about the distribution of target deviant tones. The factor SOA tested whether the presentation rate matters in extracting stimulus statistics (Tavano et al., 2014; Wacongne et al., 2012). In a first session (*uninformed*), participants were asked to detect the low pitch tone (target) with tones delivered first at a *slow* rate (constant SOA = 750 ms) and then at a *fast* rate (constant SOA = 250 ms). In a second session (*informed*), participants were explicitly informed about the repeating sequence structure and target probability density with each sequence and asked to use this information to detect targets at slow and fast stimulus rates. In sum, each participant received four orderly conditions: 1) *slow uninformed*; 2) *fast uninformed*; 3) *slow informed*; 4) *fast informed*.

### Behavioral results

Accuracy in target detection was high and did not differ among conditions (all *Fs*(_1,25_) ≤ 2.82, all *ps* ≥ 0.12; Figure 2A). We indexed the hazard rate distribution by fitting robust, nonparametric Theil-Sen estimators (Theil, 1950; Sen, 1968) to single-trial response times using successive potential target positions that is, elapsed time as a predictor. This resulted in one intercept and one regression coefficient (slope) per participant and condition. Intercept estimates reflect condition-wise changes in response speed magnitude unrelated to elapsed time. The analysis of intercepts showed that participants were quicker at responding to targets appearing within faster rather than slower sequences (*F*_(1,25)_ = 40.25, *p* < 0.001; Figure 2B) but there was no effect of prior information and no interaction (all *Fs*_(1,25)_ ≤ 1.12, all *ps* ≥ 0.30). The average response speed for fast sequences was 348 ms (Standard Error, SE = 22), and 411 ms for slow sequences (SE = 36).

**Figure 2.**
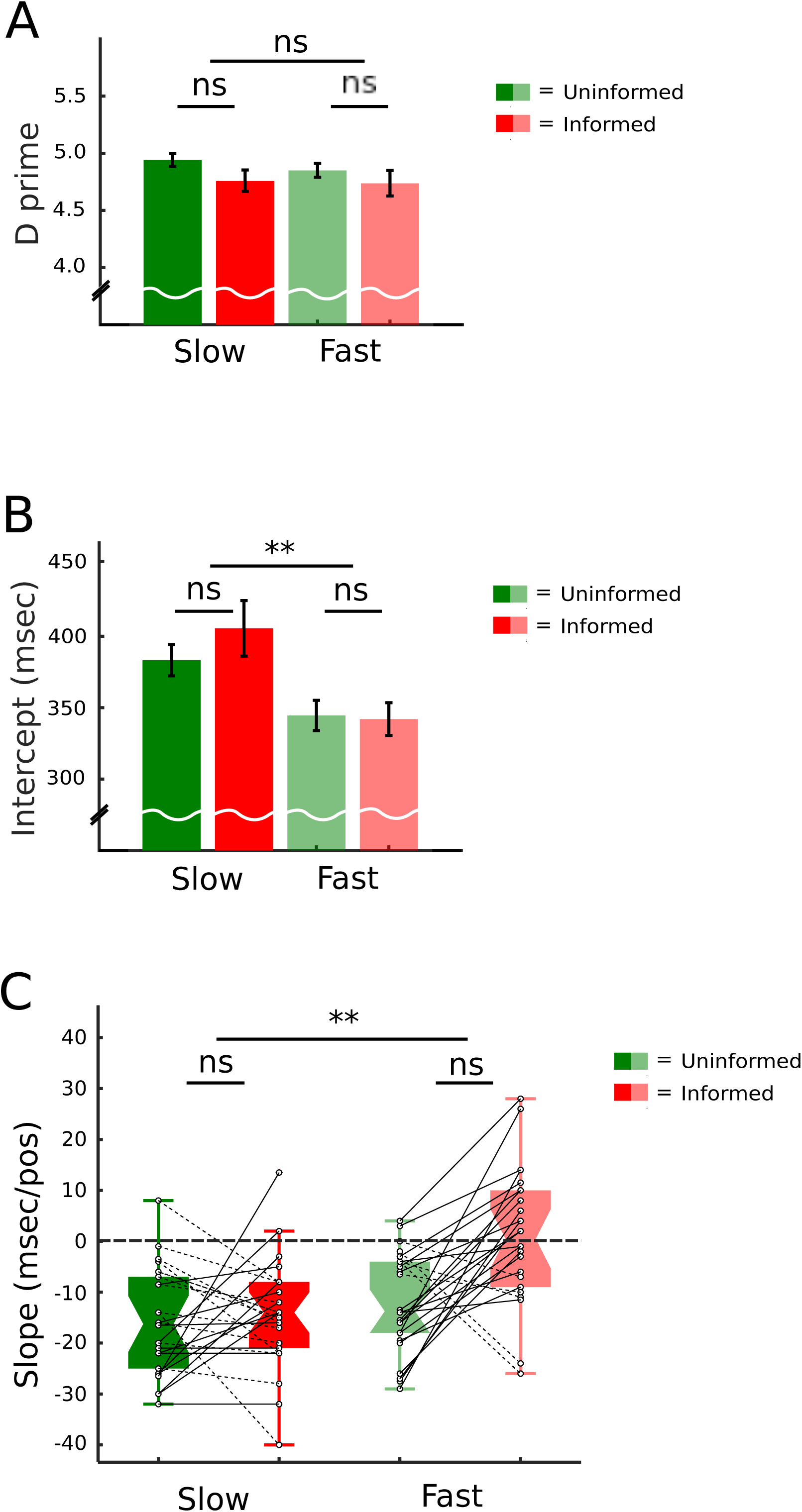
Behavioral results. A: Target detection sensitivity measures (collapsed across potential target position). Error bars represent the Standard Error of the Mean (SEM). B: Means ± SEM of estimated response time intercepts across potential target positions, showing no statistically significant difference among conditions. C: Boxplot with median (middle of each box), interquartile range and whiskers (1.5 time the interquartile range) of estimated response time slopes suggesting the presence of the hazard rate driving response speed in slow stimulus trains, regardless of prior knowledge (participants being uninformed or informed about actual target probabilities), and in fast stimulus trains with no prior knowledge (participants being uninformed). Prior knowledge in fast stimulus trains effectively cancels time-dependent (position-wise) changes in response time, that is, the hazard rate.

Theil-Sen slopes convey changes in speed using a signed single value (negative = hazard rate, as response latencies decrease with elapsed time). The distribution of Theil-Sen slopes was normal in all conditions (all Shapiro-Wilks Statistics ≥ 0.84, all *ps* > 0.21). We found a significant *Prior information* × *SOA* interaction (*F*(_1,25_) = 4.73, *p* < 0.05), driven by a significant effect of prior information in fast sequences (*t*_(1,25)_ = −3.33, *p* < 0.01). A robust hazard rate effect was present in the fast uninformed condition (*t*_(1,25)_ = 0.05, *p* = 0.95); however, the hazard rate was cancelled in the fast informed condition (*t*_(1,25)_ = 0.05, *p* = 0.95). There was no difference in the intercepts for fast sequences. A robust hazard rate was also found in both slow conditions regardless of information status (all *ts*_(1,25)_ ≤ −6.06, all *ps* < 0.001). Overall, the hazard rate effect resulted in a speed gain of 10-15 ms per potential target position (Figure 2C and Supplemental information, Figure S1A).To verify whether prior information in fast sequences reflected individually consistent cognitive changes rather than the cancelling out of random behavioral patterns at the group level, we subtracted informed from uninformed slope estimates at either SOA level and tested the observed data against the null hypothesis that prior information is equally likely to suppress or enhance the hazard rate. A binomial test indicated that in slow sequences under prior information the proportion of suppression of .46 was similar to the expected .50 (*p* = 0.84, 2-sided). In fast sequences, however, prior information suppressed the hazard rate in 20 out 26 participants, that is, in 77% of our sample (p < 0.01, Figure 2C for individual goodness of fit models; Supplemental information, Figure S1B).

### EEG results

#### Hazard rate of sensory processing

Participants’ attentive searchlight was effectively deployed on individual five-tone sequences as the prediction error generated by the fifth, non-target tone reliably indexed sequence boundaries in all conditions (all *ps* < 0.05, Supplemental Information, Figure S2A).

We then analyzed the components of event-related potentials reflecting prediction error. A singletrial Theil-Sen regression was run for each time sample at electrode level (epoch duration: 750 ms, including prestimulus time from −250 to 0) and repeated for each participant and condition. Two components arose: event-related regressions coefficients (ERRCs), encoding the effects of elapsed time on event-related potentials and event-related intercepts (ERIs), encoding brain activity unrelated to the passing of time (Hauk et al., 2006).

We ran a hypothesis-free, cluster permutation analysis (Maris and Oostenveld, 2007) to determine significant activity condition-wise (relative to noise floor) and then analyzed the effect of prior information by contrasting informed and uninformed ERRC epochs within each SOA level. ERIs showed significant characteristic peaks of activity labelled as CNV (Contingent Negative Variation; Hultin et al., 1996), N1, N2b, and P3 in all conditions (all *ps* < 0.001). Stimulus rate did not substantially affect background target-related neural activity. Prior information modulated only the late, attentive components of prediction error (N2b and P3, Supplemental Information, Figure S2B; Chennu et al., 2013).

As for ERRCs in fast sequences, we found a significant increase in brain activity in the N1 and P3 ranges, with a characteristic topography (Näätänen and Picton, 1987; Nieuwenhuis et al., 2011) in the uninformed condition (p < 0.001; Figure 3A, upper panel), while only a significant late negativity cluster was found in the informed condition (> 410 ms post onset, p = 0.05). When testing the effects of prior information, two significant clusters emerged again at N1 and P3 latencies (Figure 3A, lower panel). A negative cluster (larger N1 in the uninformed condition) began very early after target tone onset (cluster latency 58-156 ms, *p* = 0.018). A positive cluster (larger P3 in the uninformed condition) was found at 257-322 ms latencies (*p* = 0.016). Notably, deviant N1 amplitudes significantly predicted response time slopes regardless of information status (uninformed, *rho* = 0.51, *p* < 0.01; informed, *rho* = 0.61, *p* = 0.001, Steiger’s *Z* = −0.52, *p* > 0.60, Figure 3B, upper panel), while P3 amplitudes predicted behavior only in the informed condition (uninformed, *rho* = 0.01, *p* = 0.99; informed, *rho* = 0.43, *p* < 0.05, Figure 3B, lower panel). This suggests that although prior information suppressed the amplitude of both early and late prediction errors, it did not reduce but, rather, enhanced the precision of perceptual encoding relevant for behavior, suggesting it brought about consistent suppression effects at an individual level. Indeed, we verified this conclusion by calculating the correlation between N1 and at P3 activity at each scalp electrode for either information condition. A resampling analysis showed that only in the informed condition did deviant N1 activity significantly predict P3 activity, both being suppressed (*p* < 0.001; Supplemental Information, Figure S2C). The suppression effects were different at early sensory and late attentive stages. A topographical analysis (TANOVA; Murray et al., 2008) showed that prior information at P3 latency attenuated activity in the same generators or generators with a similar configuration, while it changed the generator configuration in the N1 range, between 55 and 100 ms post-onset (Supplemental Information, Figure S2D). We conclude that in fast sequences, the effects of elapsed time are such that enhancing neural afferent activity (N1 wave, Budd et al., 1998) increases response speed, and, correlatively, suppressing it cancels any speed gain.

**Figure 3.**
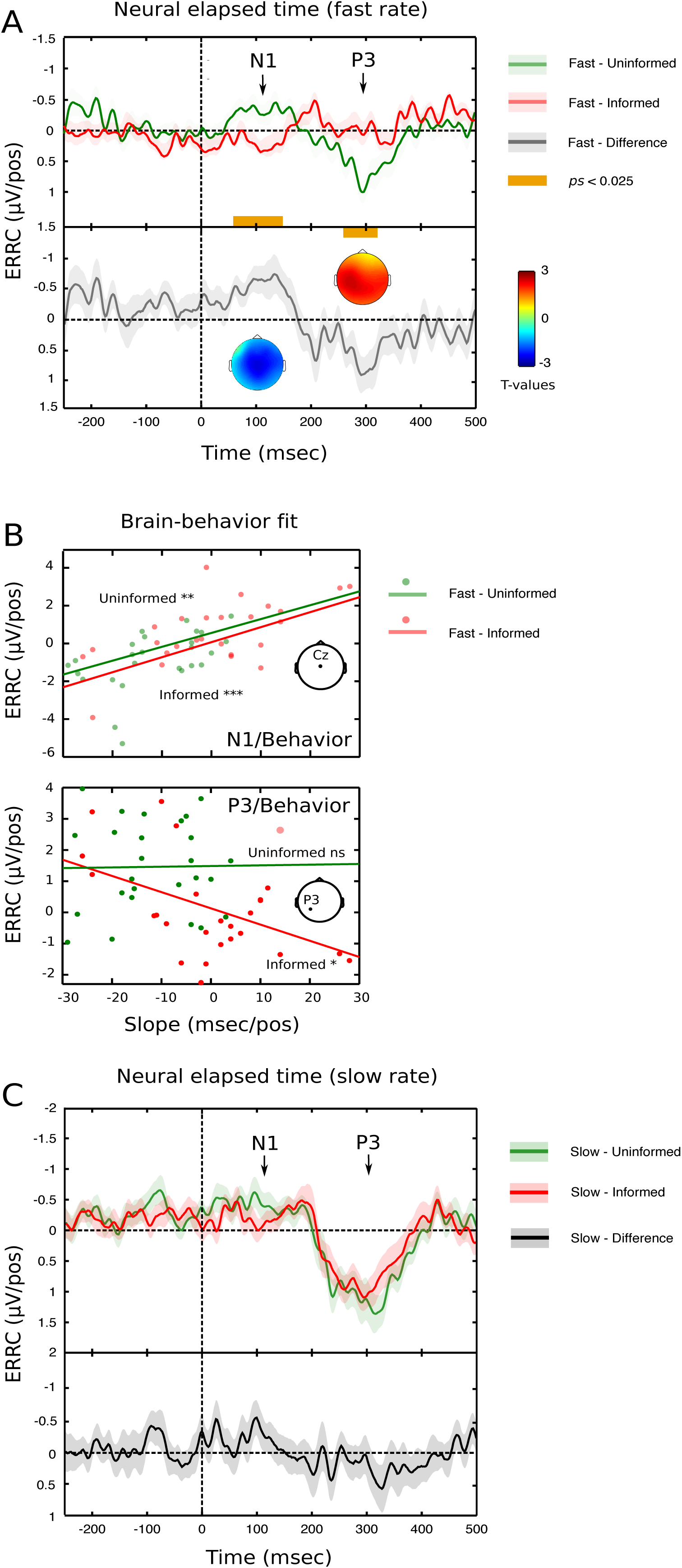
A. The hazard rate of sensory processing was imaged using Event-related Regression Coefficients (ERRCs) in fast sequences. In the uninformed condition, significant deflections in ERRC activity were found at N1 and P3 latencies. There was no significant activity in the informed condition for the first ~400 ms after target onset. ERRC cluster statistic values averaged across the electrode space show a significant effect of Prior information at N1 and P3 latencies, with a central distribution for the negative cluster and a left-sided centroparietal distribution for the positive cluster, consistent with the N1 and P3 interpretations. B. ERRCs at N1 latency in fast sequences significantly predict behavior regardless of prior information; this link is preserved for later, attentive processing (P3) in the informed condition. C. ERRCs in slow sequences show no statistically significant effect of Prior information: however, significant ERRC activity at N1 in the uninformed condition, and at P3 latencies in either information condition, highlights a hazard rate effect of sensory/attentive processing.

ERRCs in slow sequences showed significant brain activity in the P3 range regardless of information status (all *ps* < 0.001; see Figure 3C). The N1 deflection was significant only in the uninformed condition (*p* < 0.01, informed: p = 0.28). Interestingly, in either information condition N1 amplitudes better predicted response times (uninformed, *rho* = 0.57, *p* < 0.01; informed, *rho* = 0.52, *p* < 0.01) than P3 amplitudes (uninformed/informed, *rho* < 0.2, *p* > 0.10). This confirms the link between early sensory processing and response times at an individual level regardless of prior information. We found no significant effect of prior information in slow sequences. A fast stimulation may be better suited at automatically extracting and maintaining the statistical regularities of stimulus sequences on which prior information operates (Bendixen et al., 2008).

#### Hazard rate of pre-stimulus oscillations

The time domain analysis allowed detecting only post-stimulus effects. However, our main hypothesis was that brain activity should reflect endogenous probability estimates based on potential target onsets and thus predict the response time to a target before its eventual onset. In particular, for participants who were uninformed about target statistics we expected pre-stimulus single-trial power in the Beta 1 range (< 20 Hz) to increase with elapsed time, encoding the task relevance of a potential target onset. To test this we imaged the rapid update of perceived target probability by measuring the power spectrum of each trial epoch, from 5 to 28 Hz. The Theil-Sen estimator was applied to each time-frequency bin, obtaining a distribution of signed time-frequency regression coefficients (TFRCs), measured as μV^2^ unit change per potential target position with negative/positive sign equaling time-dependent decrease/increase. Unlike ERRCs, which are akin to event-related potentials, TFRCs lack clear morphological markers of neural activity. Therefore, to map out which part of the time-frequency spectrum was predictive of behavior, we ran a rank correlation analysis between each TFRC bin and the corresponding slope of response times. Such an analysis allows isolating the components of spectral power change that are task relevant. Notice that, although our hypothesis concentrated on the pre-stimulus period, the correlation analysis was run on the entire epoch, to provide a fair chance of detecting post-stimulus modulations in alignment with time domain results (Supplemental Information, Figure S3A).

Our main hypothesis was confirmed in fast sequences. In the uninformed condition, Beta 1 power (15-19 Hz) averaged across all scalp electrodes inversely correlated with response behavior ~ 150-125 ms before a potential target onset (*p* < 0.05, Figure 4A, left panel; Figure S3A). Notice that participants could not foresee whether a target would appear or not at any potential position within the current sequence. This suggests that an increase in Beta 1 oscillations encodes a portion of the variance in the neural signal representing the update of time-dependent probability estimates for potential future events, which, in turn, drives the time-dependent reduction in response times when eventually the target does appear. The increase in Beta 1 was significant at left centroparietal electrodes (*p* < 0.05, Figure 4A, right panel). Importantly, it also predicted post-stimulus ERRC N1 amplitudes (*p* = 0.01), suggesting that pre-stimulus Beta 1 power determined the distribution of response times by changing the weights of early, post-stimulus auditory processing.

**Figure 4.**
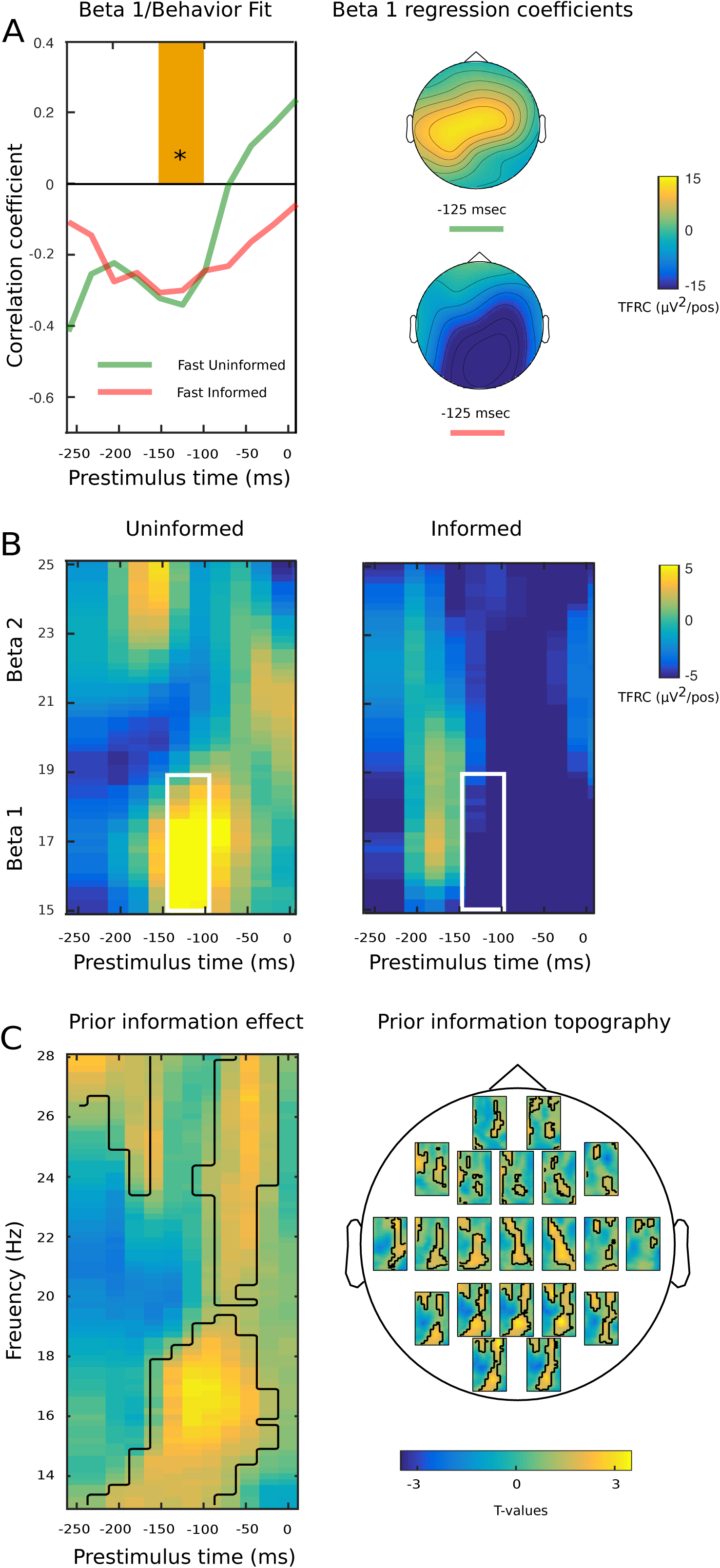
A. Left panel, prestimulus Beta 1 (14-19 Hz) oscillatory power (median across all scalp electrodes) significantly predicts response speed to eventual target onset in fast sequences, similarly for uninformed and informed conditions. Right panel, topography of Time-Frequency Regression Coefficients (TFRCs, measured in μV^2^ per potential target position, median across Beta 1 frequencies) at the behaviorally significant prestimulus interval. In the uninformed condition, the hazard rate is reflected at central electrodes; in the informed condition, prior information about actual target onset cancels the hazard rate at central electrodes and leads to significant decrease with elapsed time at parietal electrodes. B. Grand median time-frequency representation of oscillatory power regression coefficient distribution at both Beta 1 and Beta 2 in uninformed (right panel) and informed (left panel) fast sequences. White squares indicate the time-frequency bins averaged to obtain the B1/behavior fit in 4A. C. Left panel, cluster-based significance distribution of Prior information across beta-band frequencies at Pz electrode. The center of gravity lies in the Beta 1 frequency range, between 150 and 50 ms before the eventual onset of the uncertain target: These bins are fully included in the independently identified behaviorally significant prestimulus interval. Right panel, topography of the effect of prior information.

Importantly, when participants were provided with prior information, the correlation with behavior was invariant in terms of strength, latency, direction, and frequency specificity (*p* < 0.05, average across all scalp electrodes, Figure 4A, left panel; Figure S3A). Similarly, Beta 1 power for informed participants maintained a significant correlation with post-stimulus N1 amplitudes (*p* < 0.05), confirming the inference of a causal link between pre-stimulus Beta 1 power and early auditory processing. However, prior information silenced time-dependent probability update processes in the Beta 1 band at central electrodes and determined a significant decrease in spectral power at parietal electrodes (*p* < 0.05, Figure 4A, right panel). The resulting “inverse hazard rate” at parietal generators may reflect how the brain progressively silences the hazard rate at more sensory-specific central electrodes. There was no significant correlation with behavior for either Alpha (8-12 Hz, all *ps* > 0.2) or Beta 2 (21-25 Hz, all *ps* > 0.08, Figure S3B) bands.

We again resorted to a cluster permutation approach to verify the effect of prior information on single-trial power estimates, this time jointly analyzing Beta 1 and Beta 2 (14-28 Hz) and separately the alpha band (8-12 Hz). There was no effect in the alpha band (all *ps* > 0.32). In the beta band, a significant cluster with a center of gravity at parietal electrodes in the low frequency range (cluster latency ~ −200 to −50 ms, *p* < 0.05, cluster frequency 15-19 Hz, Figure 4C) emerged, confirming the suggestion that prior information may rely on a parietal network to exerts its silencing effects on elapsed time computation at central electrodes. Slow sequences presented with predominantly post-stimulus effects (Figures S3C and S3D).

## Discussion

The hazard rate reflects how the brain uses elapsed time to dynamically update the expectations for a future deterministic target to occur (Janssen and Shadlen, 2005; Leon and Shadlen, 2003). In principle, participants could simply estimate the probability density function of a target onset over trials, and use this information to respond (Luft et al., 2015). However, “that is not the natural way one thinks about it as the [waiting] process unfolds in time. Rather, if the event has not yet occurred, one senses there is some tendency for it to occur the next instant in time” (Luce, 1986, p. 13). We tested whether such a “feeling” or mounting temporal expectations (Coull, 2009; Luce, 1986; Nobre et al., 2007) would hold also for a target event that may or may not onset within a fixed time window. We found clear behavioral evidence that this is the case: A robust hazard rate was found for stimuli delivered at both slow (1.33 Hz) and fast (4 Hz) rates. Participants naïve to the target distribution became progressively faster at responding to the eventual onset of a target across potential target positions. However, when they were informed about the equiprobable target distribution, the hazard rate was largely suppressed in fast sequences as prior information factored out elapsed time from probability estimation processes. This did not happen in slow sequences, possibly because a fast stimulus rate allowed participants to more easily represent five-tone sequences in working memory (Schulze and Tillmann, 2013). Notice that the attenuation of the hazard rate in fast sequences was not due to increased variance at the group level but rather to suppressive processes acting at the individual level (~ 77% of our sample).

When we highlighted the component of the neural activity time-courses sensitive to within-sequence elapsed time (Event-Related Regression Coefficients, ERRCs), we found an increase in early post-stimulus sensory activity in the N1 range in the uninformed condition. This increase in brain activity explains a significant proportion of individual level variance in response speed, suggesting that modulations of N1 coefficients encode time-dependent information (Bertelson, 1967). This correlative finding was verified when prior information suppressed the N1 deflection in fast sequences, while remarkably maintaining a consistent predictive relationship to response behavior. We conclude that the subjective nature of probability encoding changes the weights of early sensory processing.

A second, most relevant piece of evidence comes from the analysis of power spectrum in the pre-stimulus period. Low beta-band oscillatory power (Beta 1 = 15-19 Hz), ~150-125 ms before the potential onset of the target event, consistently predicted response speed and sensory regression coefficients in both uninformed and informed fast sequence conditions. For uninformed participants, with elapsed time factored in, Beta 1 increased at central electrodes. When elapsed time was factored out by prior information, Beta 1 did not change at central electrodes and was significant suppressed (“inverse hazard rate”) at parietal electrodes. The latter result could reflect how a distributed neural system dynamically estimating event probability silences the sensory effects of elapsed time that are behaviorally relevant. An increase in Beta 1 coefficients is in line with the original proposal of Engel and Fries (2010) as well as the findings of Arnal et al. (2015) and Fujioka et al. (2012, 2015), expanding on this work by demonstrating that power modulation in the beta band is not simply relevant for temporal tracking or active sensing of event onset, but reflects internal estimates of event probability in time, which change depending on contextual information. In line with this, these results also contribute to the proposal put forward by Spitzer and Haegens (2017) who suggest a primary role of Beta 1 rhythm in the endogenous re-activation of task-relevant information (see also Bressler and Richter, 2015; Donner et al., 2009; Haegens et al., 2011; Lee et al., 2013, Michelareas et al., 2016; Stanley et al., 2016). A potentially fruitful endeavor for future research would be to bridge human and animal research by testing the relationship between cortical Beta 1 power and the encoding of the hazard rate at the neuronal level (Janssen and Shadlen, 2005; Leon and Shadlen, 2003; Yang and Shadlen, 2007).Interestingly, the neural mechanisms underlying the hazard rate constitute a peculiar case of perceptual bias. Participants gain a substantial behavioral advantage from constructing subjective temporal expectations rather than relying on the evidence from objective probability density estimates. Indeed, providing truthful prior information on the target’s uniform probability density cancelled any behavioral advantage in fast sequences. Therefore, the hazard rate of events represents an interesting conceptual challenge for approaches to perception that construe the fit between brain and behavior as solely based on precise internal models of actual event statistics (Alink et al., 2010; Friston, 2005; Spratling, 2016).

Contrary to previous work with deterministic targets (Rohenkohl and Nobre, 2011; Wilsch et al., 2014), we did not find significant regression coefficient effects in the alpha band. This may be due to the sequential cycling of attention, which may have entrained the alpha band to regular slower rhythms or to the fact that we used a probabilistic target. It is also possible that the use of target deviant sounds may have accentuated low beta relative to alpha activity (see Chang et al., 2016 for beta activity following unpredictable deviants).

It has been postulated that changes in neural rhythms may underlie different cognitive operations (Kopell et al., 2010). Notably, simulations studies of cortical rhythm formation postulate that the Beta 1 rhythm reflects memory of stimulation history, such that its modulation can distinguish between standard and deviant events (Kopell et al., 2011). The importance of cortical Beta 1 in defining the task-relevance of sensory stimuli might rely on contributions from basal ganglia generator circuits (Leventhal et al., 2012). Interestingly, recent clinical work purports that beta desynchronization in Parkinson’s disease (PD), predominantly of basal ganglia origin, specifically impairs pre-stimulus beta-band activity in rhythmic auditory perception, suggesting a causal relationship with timing deficits that are typically present in PD (Gulberti et al., 2015). This could also provide a test case to better analyze the functional specificity of synchronization and desynchronization in the low vs. high beta band, contrasting movement initiation and task relevance (Joundi, et al., 2013; Pfurtscheller et al., 2003: Pogosyan et al., 2009; Swann et al., 2009).

## Conclusions

Pre-stimulus low beta, or Beta 1 rhythm, reflects how participants think ahead of time the likelihood of a future uncertain target. The contextual modulation of Beta 1 via the absence/presence of prior information does not impair its predictive value relative to response times, suggesting it flexibly encodes internal probability estimates.

## Acknowledgements

This work was supported by a Deutsche Forschungsgemeinschaft (DFG, German Research Foundation) Reinhart-Koselleck Project grant awarded to E. Schröger and by a DFG grant (2268/6-1) awarded to Sonja A. Kotz. The authors would like to thank Matt Craddock, Maren Grigutch, Philipp Ruhnau, Michael Schwartze, Burkhard Maess, Peter Lakatos, and David Poeppel for insightful discussions and help with the experimental setup and data analyses, and Katrin Ina Koch for data collection.

## Author Contributions

A.T. devised research question; A.T., E.S., and S.K. defined research design and data collection; A.T. analyzed data and created illustrations; A.T. and S.K. wrote a draft manuscript; A.T., E.S., and S.K. revised and approved the final manuscript.

## STAR methods

### Participants

The experiment was conducted at the Max Planck Institute for Human and Cognitive Brain Sciences, Leipzig (Germany). Thirty healthy young adult individuals (15 female; age range = 19-31, mean = 25, SD = 3.5) were recruited from the institute’s database of participants. All individuals had university-level education and were paid for their participation. Four participants were excluded from further analysis: two for below-average behavioral performance (less than 50% target detection), one for misinterpreting task instructions, one for excessive target rejection rate after Independent Component Analysis (pass cutoff: 80%, i.e. at least 16 target trials per position and condition). The final pool of 26 participants (13 females) reported no neurological or psychiatric disorders or therapies involving the central nervous system. Individually measured, bilateral audiometric thresholds of at least 30 dB Hearing Level at 0.25 - 8 KHz octave frequencies (ANSI, 1996). All participants signed a written informed consent complying with the Declaration of Helsinki on human experimentation, and approved by the Ethics Committee of the University of Leipzig.

### Stimuli

Stimuli were three 50-ms pure tones (5 ms rise/fall), binaurally presented via loudspeakers at ~80 dB SPL and generated using Matlab (version 7, Mathworks, Natick, MA). A 440 Hz tone (A4 on the equal tempered scale), termed *standard*, was presented 900 times per condition (75% global probability). A 494 Hz tone (B4, two semitones higher than the standard), termed *non-target deviant*, was presented 240 times per condition (20% global probability). A 349 Hz tone (F4, four semitones lower than the standard), termed *target deviant*, was presented 60 times per condition (5% global probability). Stimuli were delivered using Presentation© software (version 12.0, www.neurobs.com) running on a Windows PC.

### Experimental Design

Participants sat in an electrically shielded, sound-attenuated chamber, and fixated a white cross on black computer screen at a distance of ~1 meter while listening to the auditory stimuli. They responded to target tone onset by pressing a button on an external response box. Figure 1 illustrates the experimental design. Standard (440 Hz, A_4_) and non-target deviant tones (496 Hz, B_4_, two-semitone difference relative to standard) were organized as continuously repeating sequences of four standards followed by a non-target deviant in fifth position, without interruptions. Target deviant tones (349 Hz, F_4_, four-semitone difference relative to standard) appeared rarely (20% of sequences) and unpredictably at either standard position two, three or four, with a uniform distribution (20 targets per position). The distribution of target-containing sequences was individually randomized for each condition and participant, with two constraints: 1) a maximum of one target per sequence, 2) a minimum of two sequences without targets between two target-containing sequences.

Targets were equiprobably distributed across standard position two, three and four, yielding a discrete uniform distribution function: f(t) = 1/3, for each of standard positions two, three, and four. Denoting the survival probability (“the event has not yet occurred”) as 1 − F(t), where F(t) is the cumulative distribution function, the hazard function is then: h(t) = f(t)/(1-F(t)). See Figure B.The experiment was organized into two parts, with a fixed order. In the first part, participants were instructed to respond to the onset of target tones as accurately and fast as possible by pressing a button on a response box. They trained in a short block of 60 experimental randomly distributed tone sequences containing three targets. If errors were made (Missing, False Alarm), the training block was repeated until no errors were detected. Experimental tone sequences were first delivered with a constant 750-ms stimulus onset asynchrony (SOA), corresponding to 1.6 Hz stimulus rate (three 5-min blocks; first condition), and then – after a short break – with a constant 250-ms SOA, corresponding to 4 Hz stimulus rate (one 5-min block; second condition). The first two conditions were tagged as *slow uninformed* and *fast uninformed*, respectively.

In the second part, after a 15-min break, participants were informed, both verbally and using visual aids, about the structure of the repeating tone sequence and target probability distribution statistics within a sequence. The instruction therefore changed: they were asked to respond to target deviant tones whose onset would break the sequence at either standard position two, three, or four, with equal probability. They again trained with a short block of 60 tones and 3 targets. If errors were made (Missing, False Alarm), the training block was repeated until no errors were detected. The experimental tone trains were again first delivered with a 750-ms constant SOA (three 5-min blocks; third condition), and then – after a short break – with a 250-ms constant SOA (one 5-min block; fourth condition). These conditions were tagged *slow informed* and *fast informed*, respectively. All participants completed the experiment in about one hour.

### Behavioral Data Analysis

Hits were all button presses whose response time ranged between 150 and 1250 ms from the onset of a target deviant tone. Button presses recorded after 1250 ms were considered as false alarms (FA). Accuracy was measured by z-transforming hit and false alarm counts (5% adjustment for ceiling effects) to obtain a d’ index of task sensitivity (zFA – zHit: Macmillan and Creelman, 1991). As for the hazard rate of response times, we assumed that over three time points the true trend is effectively linear. A linear fit allowed estimating at the same time the presence of the hazard rate (negative slope = shorter response times for longer waited for events) and its strength (slope magnitude). Single-trial Theil-Sen estimates of individual linear fits across the three target positions were obtained. The Theil-Sen estimator represents an unbiased, robust, non-parametric simple linear regression method, which extracts the median slope among all possible pairwise combinations of points (Theil, 1950; Sen, 1968). Accuracy and reaction time data entered a twoway, repeated-measures ANOVA, with the factors SOA (slow, fast) and prior information (uninformed, informed). Results with p ≤ 0.05 were declared significant. Greenhaus-Geisser correction was applied whenever Mauchly’s test signaled violation of the sphericity assumption.

### EEG Data Acquisition and Preprocessing

Electroencephalographic (EEG) data were collected using a 26 scalp Ag/AgCl electrode set (BrainAmp), mounted in an elastic cap according to the 10-20 International system. The electrode space was composed by: Fp1, Fpz, Fp2, F7, F3, Fz, F4, F8, FT7, FC3, FC4, FT8, T7, C3, Cz, C4, T8, CP5, CP6, P7, P3, Pz, P4, P8, O1, O2. Two external electrodes were placed at right and left mastoid sites. For electrooculographic (EOG) data recording, four additional electrodes were placed at both eye canthi, and above and below the right eye. For participants 19 to 26 (21-30 in the original dataset), the cap contained 38 more electrodes (10-10 system), not used in the current analysis for comparability across participants (Fpz was excluded from participants 1-18 as it was conversely not present in participants 19-26). An online reference was placed on the tip of the nose and the sternum served as ground. Electrode impedance was kept below 5 kQ. EEG/EOG sampling rate was set to 500 Hz, with online highpass filtering at 0.01 Hz. The resulting continuous recordings were visually inspected and pruned from non-stereotypical artifacts or extreme voltage changes values. An Independent Component Analysis (ICA, extended Infomax, Delorme and Makeig, 2004) was performed on the pruned continuous data, offline bandpass filtered 1-100 Hz (Kaiser window, Beta 5.6533, filter order 1812 points, transition bandwidth 1 Hz, see Widmann et al., 2014). The maps of exemplar Independent Components (ICs) reflecting blinks or vertical eye movements and horizontal eye movements from one participant were selected as spatial templates in a semi-automatic artifact search across all ICs of the remaining datasets (correlation threshold, *r* = 0.94, Viola et al., 2009). Eye-movement-related ICs, both vertical/blink-related and horizontal, ranged between 1 and 3 per participant. Selected ICs were verified in their spectral power distribution before being subtracted (Onton and Makeig, 2006). The resulting continuous datasets were finally low-pass filtered at 35 Hz (filter order 184, transition bandwidth 10 Hz).

### Event-related potential and coefficient analysis

Epochs were separately extracted for the onset of standard, non-target deviant and target deviant stimuli, beginning 1000 ms before and ending 1000 ms after stimulus onset. Epochs were selected based on their effective contribution to increasing the signal-to-noise ratio (Rahne et al., 2009). On average, 12.3% of epochs were rejected. Epochs of interest for regression analysis began 250 ms before the onset of target deviant tone, and ended 500 ms thereafter. Prediction error responses for the non-target deviant trials were calculated in sequences that did not contain a target trial by subtracting the average response of the fourth standard tone. As for target trials, a minimum of 16 trails (80%) per Target position and conditions was retained. For each time point of each trial, a Theil-Sen estimate of the linear relationship between target position – a proxy for elapse time – as predictor and event-related electrical activity was calculated. There resulted event-related regression coefficients (ERRCs, slopes) and event-related intercepts (ERIs). As stimulation was isochronous, target position directly reflects elapsed time. Therefore, ERRCs encode neural estimates of elapsed time (Dien et al., 2003; Hauk et al., 2006). We took a data-driven approach to analyze the distribution and polarity of neural effects within a post-stimulus window of interest (0-600 ms). The presence of differences in the effects of Target position (elapsed time) as determined by Prior Information was tested for all channels/time points separately at each SOA level: slow uninformed vs. slow informed; fast uninformed vs. fast informed. ERRCs entered a non-parametric, cluster-based permutation test of significance, which allows controlling for type I error rate in the presence of multiple comparisons (Maris and Oostenfeld, 2007). Clusters were minimally composed of two electrodes. Cluster-level statistics was determined by summating significant paired T-test values (cluster alpha = 0.05) across adjacent points within each cluster, and evaluated under the distribution obtained by drawing 1000 within-subject, random permutations of the observed data. Noise floor was estimated by randomizing individual data points along the time axis. Results show the probability (alpha = 0.05) of obtaining a cluster-level statistic that is larger (positive polarity) or smaller (negative polarity) than the observed one. Topographical differences in the distribution of current density were investigated using the Global Dissimilarity Index, which measures the configuration of electric fields (and their linear transformations), normalized by their individual strength Global Field Power (Murray et al., 2008).

### Time-frequency analysis

To increase signal-to noise ratio, data were subject to a Principal Component Analysis, and the first four components, accounting on average for 93% of variance, were retained for further processing. Zero-padded (5 s), individual target epochs were submitted to a time-frequency analysis at each electrode using a Morlet wavelet (7 cycles, estimated for the central ± 4 SD of the Gaussian envelope, Oostenveld et al., 2011), for frequencies comprised between 5 and 28 Hz, in steps of 0.25 Hz. Event-related power estimates of target position effects at each frequency/time point were extracted from 500 ms pre-stimulus to 500 ms post-stimulus (sliding window = 25 ms). Each time-frequency bin entered a Theil-Sen, non-parametric regression analysis with Target position as predictor, obtaining time-frequency regression coefficients (TFRCs), and time-frequency intercepts (TFIs). We obtained median TFRC power estimates across all electrodes at Alpha (8-12 Hz), low beta (Beta 1, 14-19 Hz) and Beta 2 (20-25 Hz) rhythms, and calculated a Kendall-type (Kendall, 1938) rank correlation between each TFRC and response time slope, in order to determine the time-frequency band relevant for behavior. A permutation resampling approach (1000 repetitions) was used to test the significance of rank correlations.

To analyze the effect of Prior information within each SOA level, we resorted to a cluster-based permutation approach of power estimates between 250 ms pre-stimulus and 500 ms post-stimulus, using 2-tailed T-tests (significance set at 0.05), which controls for the multiple comparisons problem (all scalp electrodes, minimal N electrodes per cluster = 1, T-test cluster parameter = maxsize, 1000 permutations, cluster alpha set at 0.1; Maris and Oostenveld, 2007). All analyses were run using EEGLAB (Delorme and Makeig, 2004), FieldTrip (Oostenveld et al., 2011), and custom Matlab scripts.

## Supplemental Information

Supplemental information includes three figures, illustrating raw response times and goodness-of-fit results, a proof of attention deployment on five-tone sequences (attention reset role of non-target deviant), the predictive relationship between N1 and P3 activity in fast sequences, and the time-frequency analysis of Target responses in fast and slow sequences, across the whole epoch.

